# Complex rheology of condensin in entangled DNA

**DOI:** 10.1101/2025.06.04.657969

**Authors:** Filippo Conforto, Antonio Valdes, Willem Vanderlinden, Davide Michieletto

## Abstract

Structural-Maintenance-of-Chromosome (SMC) complexes such as condensins are well-known to dictate the folding and entanglement of interphase and mitotic chromosomes. However, their role in modulating the rheology and viscoelasticity of entangled DNA is not fully understood. In this work, we discover that physiological concentrations of yeast condensin increase both the effective viscosity and elasticity of dense solutions of *λ*-DNA even in absence of ATP. By combining biochemical assays and single-molecule imaging, we discover that yeast condensin can proficiently bind double-stranded DNA through its hinge domain, in addition to its heads. We further discover that presence of ATP fluidifies the entangled solution possibly by activating loop extrusion. Finally, we show that the observed rheology can be understood by modelling SMCs as transient crosslinkers in bottle-brush-like entangled polymers. Our findings help us to understand how SMCs affect the rheology and dynamics of the genome.

---

Among the most important processes orchestrating chromosome folding in both interphase and mitosis is the formation of loops, performed by structural-maintenanice-of-chromsome (SMC) complexes, such as cohesin, condensin and SMC5/6 [1–11]. Although these complexes perform loop extrusion *in vitro* [12–14], the extent to which loop extrusion affects genome organisation *in vivo* is poorly understood [15–18].

Alternative models to loop extrusion are also able to explain some of the experimental observations, both *in vivo* and *in vitro*. For instance, bridging-induced phase separation (BIPS) can recapitulate the formation of clusters, or condensates, of yeast cohesin in the presence of DNA [19]. At the same time, the loop capture model can explain the topological trapping of a DNA plasmid by a condensin that is loaded on a tethered linear DNA [20]. Both these models rely on the fact that SMCs can “bridge” inter-chromosomal DNA, i.e. can simultaneously bind two dsDNA segments that do not belong to the same DNA molecule. On the contrary, intermolecular bridging is not envisaged in loop extrusion models, as the SMCs are thought to “reel in” DNA in cis [16]. Mixed models, whereby SMCs perform an *effective* loop extrusion by bridging DNA segments have also been successful at capturing some puzzling evidence, for instance the formation of Z-loops and the bypassing of large obstacles bound to DNA [21], or the observation that condensin can make steps larger than its own size [22].

SMCs can also have an impact on the dynamics of chromosomes in cells, although it is challenging to precisely quantify it experimentally [23]. The prediction from most computational and theoretical works is that loop extrusion performed by SMCs will compact [24, 25], segregate [26], fluidify [27] and even unknot [28, 29] chromatin.

However, to the best of our knowledge, there are no experimental works to verify these predictions as most of the *i* n vitro assays have so far focussed on dilute and/or tethered DNA conditions. To understand the role of SMCs *in vivo*, we must study their behaviour in physiologically dense DNA solutions. For this reason, in this work we perform microrheology [30, 31] of entangled solutions of *λ*-DNA under the action wild type (WT) and mutated (Q) yeast condensin.

Contrary to what expected from existing computational and theoretical work, we observe that condensin *increases* the viscoelasticity of entangled solutions of *λ*-DNA. Importantly, we also discover that this “thickening” is loop extrusion independent as we observe it in a catalytically dead protein complex. To better understand our observations, we then performed single molecule (AFM) and bulk measurements and discovered that the hinge – the domain through which the SMC proteins dimerize – has a significant binding affinity to dsDNA, working as a “DNA bridging patch”. We conclude our paper by performing coarse grained MD simulations of a dense solution of entangled polymers under the effect of loop extruders with additional DNA-bridging patches. We observe that in the presence of sticky DNA-bridging patches, our simulations display thickening, as observed in microrheology.

In line with the DNA bridging [19] and loop capture [18] models, our experiments suggest that condensin can bind two different dsDNA molecules *in trans* and thus transiently cross-link DNA in dense conditions. Our results contribute to understanding the action of SMC proteins in physiologically crowded and entangled conditions.

## MATERIALS AND METHODS

### Protein expression and purification

Wild-type (WT) and Q-loop condensin holocomplexes were expressed from two 2*µ*-based high copy plasmids transformed into *S. cerevisiae*. Purification of holocomplexes has been performed as in Ref. [32] (see SI for more details). The hinge domain was expressed by transforming DNA fragments encoding for yeast Smc2 residues 396-792 and yeast Smc4 residues 555-951 in bacteria. Specifically, we co-expressed Smc2 (396-792) with an N-terminal (His)6-TEV-tag and Smc4 (555-951) without a tag. Purification was done via Ni-columnn (see SI for full details).

### Electrophoretic mobility shift assay (EMSA)

The 6-FAM labeled 50-bp dsDNA was prepared by annealing two complementary DNA oligos (Merck, 5’-6-F AM-GGATACGTAACAACGCTTATGCATCGCCGCCGCTACATCCCTGAGCTGAC-3’; 5’-GTCAGCTCA GGGATGTAGCGGCGGCGATGCATAAGCGTTGTTACGTATCC-3’) in annealing buffer (50 mM Tris-HCl pH 7.5, 50 mM NaCl) at a concentration of 500 mM in a temperature gradient of 0.1 C/s from 95^*°*^C to 4^*°*^C. The EMSA reaction was prepared with a constant DNA concentration of 10 nM and the indicated concentrations of purified protein in binding buffer (50 mM Tris-HCl pH 7.5, 50 mM KCl, 125 mM NaCl, 5mM MgCl2, 5% Glycerol, 1 mM DTT). After 10 min incubation on ice, free DNA and DNA-protein complexes were resolved by electrophoresis for 1.5 hr at 4 V/cm, on 0.75% (w/v) TAE-agarose gels at 4^*°*^C. 6-FAM labeled dsDNA was detected directly on a Typhoon FLA 9,500 scanner (GE Health-care) with excitation at 473 nm with LPB (510LP) filter setting.

### Fluorescence Polarisation

Fluorescence polarization (FP) experiment was performed by mixing 20 nM of the 6-FAM labeled 50 bp dsDNA (see Methods EMSA) with series of protein concentrations, ranging from 0.03125 *µ*M to 32 *µ*M, in FP buffer (25 mM Tris-HCl pH 7.5, 100 mM NaCl, 5 mM MgCl2, 1 mM DTT, 0.05% Tween20, 0.05 mg/ml BSA). The mix was incubated for 30 min at room temperature in order to attain equilibrium. Immediately thereafter, fluorescence polarization was recorded using 485 nm and 520 nm excitation and emission filter on a Tecan SPARK Microplate reader. The change in fluorescence polarization was then plotted as mean values of three independent replicates and the dissociation constant determined.

### AFM imaging

Atomic Force Microscopy (AFM) was performed on poly-L-lysine-coated mica [33]. Linear dsDNA of 500 bp was generated by PCR from pUC19 plasmid using primers 5’-AGAGCAACTCGGTCGCCGCATA (forward) and 5’-GCTTACCATCTGGCCCCAGTGC (reverse)). We mixed 0.5 ng/*µ*L DNA and 10 nM WT condensin in aqueous buffer (50 mM Tris-HCl pH = 7.5, 25 mM NaCl, 5 mM MgCl2, 1mM DTT, 1mM ATP) and incubated at room temperature for 15 seconds before deposition. Deposition of the sample onto poly-L-lysine coated mica was done by dropcasting. After surface adsorption for 15 s, the sample was rinsed using milliQ water (20 mL) and subsequently dried using a gentle stream of filtered N2 gas. For imaging the sample we used a Nanowizard 4 XP AFM (JPK, Berlin, Germany) in tapping mode; image processing was done using MountainSPIP software (see SI).

### Microrheology

For microrheology experiments, we mixed 5 *µ*l of 500 ng/*µ*l *λ*DNA with 1 *µ*l of 2 *µ*M yeast condensin (WT or Q), 1 *µ*l of 10x condensin reaction buffer (Tris-HCl Ph 7.5 500 mM, NaCl 250 mM, MgCl2 50 mM, DTT 10 mM), 1 *µ*l of 10 mM ATP and 1 *µ*l of 2 *µ*m PEGylated polystyrene beads (Polyscience). We loaded the sample into a 100 *µ*m thick sample chamber comprising a microscope slide, 100 *µ*m layer of double-sided tape and a cover slip. We recorded movies on a Nikon Eclipse Ts2 microscope with a 20x objective and Orca Flash 4.0 CMOS camera (Hamamatsu) for 2 minutes at ∼ 100 fps on a 1024×1024 field of view, resulting in about 500 tracks per condition. Particle tracking was done using trackpy and in-house code was used to process the tracks into MSD and complex modulus following Ref. [31].

### Molecular Dynamics Simulations

Entangled DNA solutions were modelled as semiflexible Kremer-Grest linear polymers [34] with *N* = 500 beads of size *σ* = 10 nm. The beads interact with each other via a truncated and shifted Lennard-Jones potential and adjacent beads are connected by FENE springs. The persistence length of the polymers is *l*_*p*_ = 5*σ* = 50 nm and the volume fraction of the solution is around 5 %. After thorough equilibration (see SI), the polymers are loaded with *N*_*SMC*_ = {5, 25} SMCs and then either let to loop extrude as in Ref. [27], or otherwise left on the loaded state to mimic conditions with no ATP. Each SMC is decorate with patches that have an attractive interaction with the DNA beads, modelled by a Morse potential with a maximum depth of 25*k*_*B*_*T*, which is comparable with the heads (and hinge) binding affinity Δ*G* ≈ − *k*_*B*_*T* log (*k*_*D*_) where *k*_*D*_ ≃ 0.1 *µ*M. The simulation is performed in LAMMPS [35] with custom-made fixes that update the position of active loop extruders (https://git.ecdf.ed.ac.uk/taplab/smc-lammps). Specifically, our loop extrusion algorithm performs a geaotry check before updating the position of the SMCs in order to preserve the topology of the system [36]. We then track the dynamics of the polymers when the SMCs are only loaded (no loops) and when allowed to make large loops via loop extrusion. In the latter case, the polymers start to resemble bottle-brushes [24, 27]. At the same time, we perform Green-Kubo calculations of the stress relaxation modulus, i.e. we compute the autocorrelation of the off-diagonal components of the stress tensor [37] (see SI for more details).

## RESULTS

### Condensin increases the viscoelasticity of entangled DNA

To understand the effect of condensin in dense solutions of entangled DNA we studied their rheology. We hypothesised that if yeast condensin performed pure loop extrusion, i.e. with only intra-molecular contacts, we would observe a significant disentanglement and fluidification of the DNA solution as predicted in the literature and as a consequence of the formation of bottle-brush-like structures [24–27, 38] (fig. 1a,b).

**FIG. 1.**
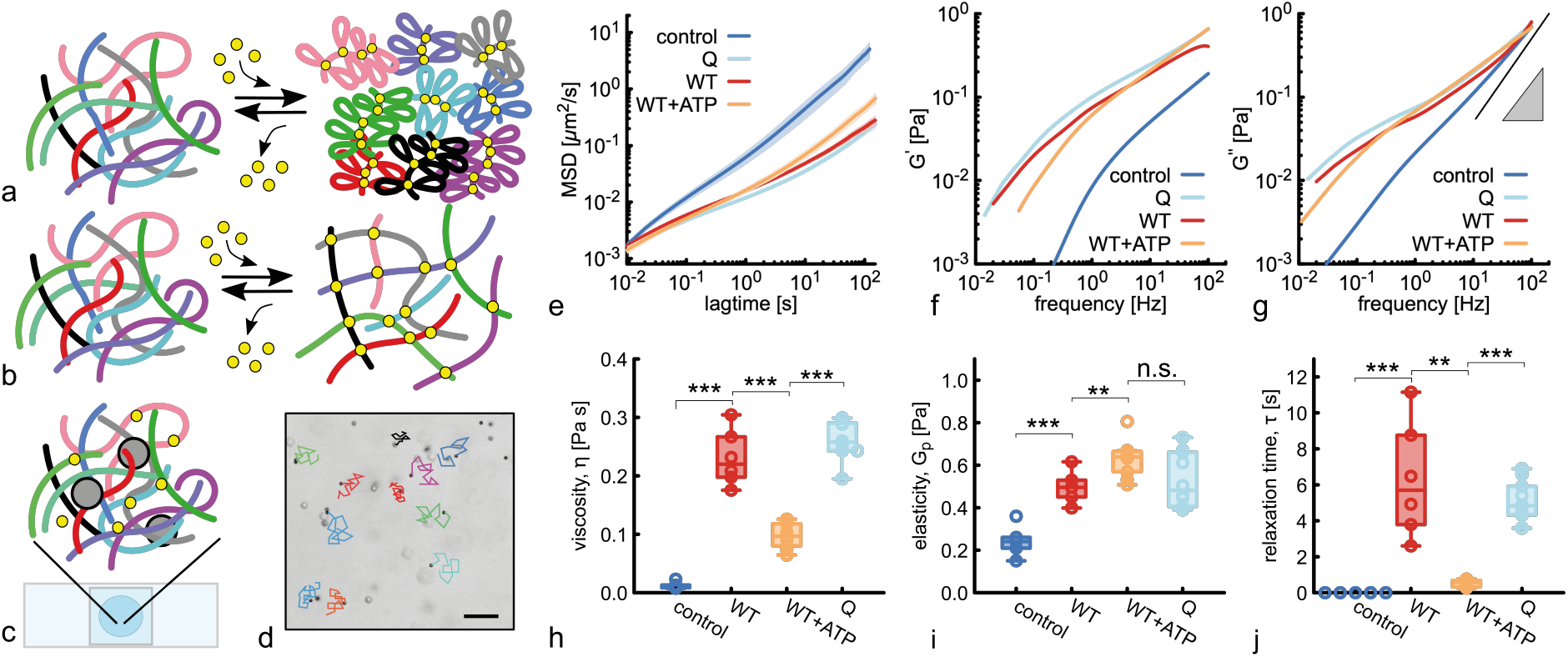
**a-b** Sketches of our two hypotheses: **a**. if condensin performed loop extrusion (intramolecular contacts only), we would expect a solution of entangled linear DNA to be converted into one made of bottle-brush-like polymers, reducing both entanglement and viscoelasticity. **b**. If condensin performed DNA-bridging (intermolecular contacts), we would expect transient crosslinks. **c**. The sample made of *λ*DNA, condensin, reaction buffer and passive tracers is mixed, incubated and then pipetted in a closed chamber. **d**. Snapshot of the field of view showing the tracers and short example trajectories (scale bar 20 *µ*m). **e**. Mean squared displacement (MSD) of the tracer beads for wild type yeast condensin (WT) in presence and absence of ATP and for a catalytically dead (Q) mutant. For all samples in this figure, DNA concentration is 250 ng/*µ*L (or 7.8 nM of *λ*DNA) and protein concentration is 0.2 *µ*M, i.e. about 25 SMCs per DNA molecule. **f-g**. Elastic (*G*^*′*^, **f**) and viscous (*G*^*′′*^, **g**) complex moduli obtained from the MSDs through the generalised Stokes Einstein relation [31]. **h**. Zero-shear viscosity, obtained from the long time behaviour of the MSD. **i**. Elasticity 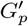 obtained from the elastic modulus measured at 100 Hz. **j**. Relaxation time *τ*_*R*_, obtained as the inverse of the smallest frequency at which *G*^*′*^ and *G*^*′′*^ intersect.

To quantitatively measure the change in rheology, we prepared samples of entangled *λ*-DNA (48’502 bp) at around 12 times the overlap concentration (*c* = 250 ng/*µ*l, *c*^*\**^ = 20 ng/*µ*l [39, 40]), and added 0.2 *µ*M of either wild type (WT) yeast condensin or a catalytically dead (Q-loop) mutant that cannot perform loop extrusion [32]. We also included 2 *µ*m-sized PEG-passivated polystyerene tracer beads and adjusted buffer conditions to that used to observe loop extrusion in single molecule assays [14, 32] (fig. 1c). We then performed microrheology, i.e. recorded videos of the passive tracers moving in the solution (fig. 1d) and extracted their mean squared displacement (MSD) *δ*^2^*r*(*t*) = ⟨[***r***(*t* +*t*_0_) −***r***(*t*_0_)]^2^⟩, where the average is performed over beads and initial times *t*_0_ (see fig. 1e).

According to most current models, SMCs should compact DNA by performing loop extrusion and thus decrease the viscosity of the entangled solution [24]. In our experiment, this fluidification would manifest itself as an increase in effective diffusion coefficient of the tracer beads and an absence of subdiffusive behaviour [40]. On the contrary, we observed the opposite: a significant decrease in the mobility of the beads and an increase in their subdiffusive regime for both WT and Q-loop condensin and even in the presence and the absence of ATP (fig. 1e). This behaviour is unexpected according to existing loop extrusion models and it points to a previously unappreciated loop-extrusion-independent DNA organisation mechanism by SMCs.

To quantify the elastic and viscous response of the fluid at different timescales we transformed the MSDs into elastic (*G*^*′*^(*ω*)) and viscous (*G*^*′′*^(*ω*)) complex moduli via the generalised Stokes Einstein relation [30, 31]. In Figs. 1f-g, one can appreciate that the presence of WT and Q-loop condensin significantly affect the shape of *G*^*′*^(*ω*) and *G*^*′′*^(*ω*). More specifically, the control displays a purely viscous behaviour with little sign of inflection in *G*^*′′*^(*ω*); on the contrary, the samples with SMCs display at least one intersection between the two complex moduli (see SI). This entails that the fluid’s response is elastic-dominated at short timescales (large *ω*) and liquid-dominated at long timescales (small *ω*). Interestingly, it is clear from the curves in Figs. 1f-g that SMC induce a significant increase in both elasticity and viscosity of the samples across all frequencies.

We compute the zero-shear viscosity of the samples *η* = *k*_*B*_*T/*(3*πaD*), where *a* = 2 *µ*m is the size of the beads and *D* the large-time diffusion coefficient obtained from the MSDs. Adding SMCs induce a 20-30x increase in viscosity in all samples, with the increase being more pronounced for WT and Q-loop mutant (fig. 1h). Interestingly, the sample with WT and ATP displays the smallest change (∼ 10 x), which may point to a fluidification effect of loop extrusion. We can also compute the large-*ω* elasticity 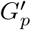 and relaxation time *τ*_*R*_ of these viscoelastic fluids by evaluating 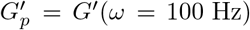 and *G*^*′*^(1*/τ*_*R*_) = *G*^*′′*^(1*/τ*_*R*_), respectively. The former (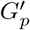, fig. 1i) suggest that the short-time elastic behaviour is significantly stiffer for SMC samples, irrespectively if with or without ATP. Despite this observation, all samples display 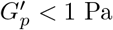, implying that they are very soft. On the other hand, the latter (*τ*_*R*_, fig. 1j) suggest that samples with WT condensin and ATP behave like liquids on a shorter timescales (smaller *τ*_*R*_) than the ones without ATP or with the Q-loop mutant which remain solid-dominated for longer times, up to tens of seconds.

Our observations suggest that SMCs may form transient cross-links between DNA molecules that are proximal in 3D space. The result is the formation of a dynamic and reversible mesh of weakly cross-linked polymers, which may be mobile in the case of loop extruding complexes. This model is in line with the “bridging-induced” phase separation behaviour observed in yeast cohesin [19] and as predicted by the loop capture model [18, 41] or modelled by the inter-molecular loop-extrusion [21]. Despite this, structural studies have only identified one set of DNA binding sites localised at the yeast condensin core (Ycs4-Brn1-SMC heads) and the anchor (Ycg1-Brn1) [32] of the complex, and we thus lack a mechanistic understanding underpinning the hypothesised transient cross-linking.

### Yeast condensin can bridge dsDNA through its hinge

To further understand the cross-linking mechanism underlying the thickening observed in the previous section, we decided to investigate different binding modes of yeast condensin to dsDNA. Condensin binds dsDNA through both its anchor domain (BrnI-YcgI) domain [32, 42] and its core subcomplex (SMC heads -Ycs4) [32], however there is no direct evidence of dsDNA binding by any other condensin domain. First, we sought computational evidence for an additional binding site by scanning through AlphaFold3 structures and found a model that predicted an interaction between the SMC2/SMC4 hinge and a dsDNA oligomer containing a small ssDNA bubble (fig. 2b). Motivated by this prediction, we performed electrophoretic mobility shift assay (EMSA) and observed a clear shift when the hinge domain (SMC2:K841-L698, SMC4:Q646-F865) was mixed with a 50 bp dsDNA segment (fig. 2c), with an estimated binding affinity of *K*_*d*_ ≃ 0.1 − 0.2 *µ*M. This measurement was further supported by fluorescence polarization (FP), where yeast condensin hinge was mixed with a fluorescently-labelled dsDNA oligo, albeit we measured a larger binding constant *K*_*d*_ ≃ 0.7 *µ*M (fig. 2d). Interestingly, these *K*_*d*_ values are comparable to – if not smaller than – the binding constants of the YcgI-BrnI (anchor) complex to DNA, i.e. *K*_*d*_ ≃ 1.7 *µ*M [42] and of the “core” subcomplex (SMC heads + Ycs4) *K*_*d*_ ≃ 0.1 − 0.2 *µ*M (see SI), both measured from *Chaetomium thermophilum*. Arguably, both EMSA and FP potentially underestimate the true *K*_*d*_ because they employ short dsDNA oligos, which are not the natural substrate for these protein complexes; however, they convincingly demonstrate that the hinge is a dsDNA binding site potentially as good as the core/anchor subcomplex.

**FIG. 2.**
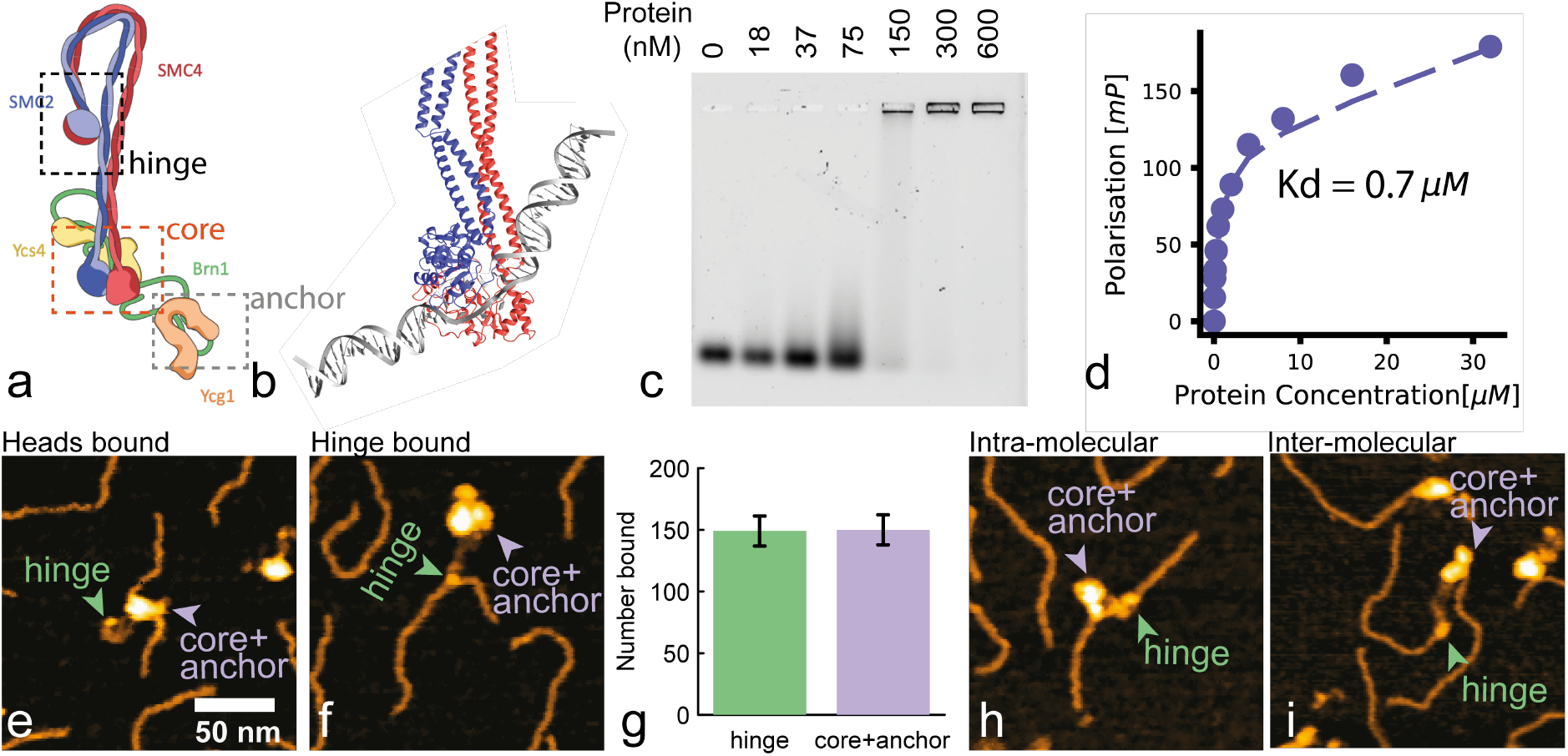
AFM characterization of WT yeast condensing complexes on DNA. **a**. Cartoon structure of yeast condensin with the hinge domain within dashed lines. **b**. AlphaFold3 model of structural interaction between the hinge domain and a segment of dsDNA with a small ssDNA bubble in the middle. **c**. EMSA showing significant binding of the hinge domain (SMC2:K841-L698, SMC4:Q646-F865) to a 25 bp dsDNA oligo in vitro with an estimated *k*_*D*_ *≃* 0.2 − 0.3 *µ*M. **d**. Fluorescence polarisation assay done with the hinge domain mixed with fluorescently-labelled 50 bp dsDNA oligo and yielding *k*_*D*_ = 0.7 *µ*M. **e-f**. Representative AFM topographs of head-bound (e) and hinge-bound (f) condensin-DNA complexes. Green and lilac arrowheads indicate hinges andheads, respectively. **g**. Quantification of relative hinge and heads bound complexes. Error bars reflect counting statistics 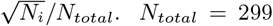. **h-i**. Representive AFM topographs of intra-molecular (h) and inter-molecular (i) condensin-DNA complexes suggesting significant “bridging” of DNA molecules *in trans*.

Motivated by these measurements, we decided to visualise dsDNA binding by the whole yeast condensin holocomplex in single molecule experiments. We mixed yeast condensin holocomplex with a 500 bp dsDNA segment, deposited it on mica and observed it using Atomic Force Microscopy (AFM, see Methods). We observed that yeast condensin displays different modes of binding: through its heads, hinge, or both (fig. 2e-f,h-i). When the holocomplex binds through its heads, we also observe a severe kink in the dsDNA molecule, in agreement with the cryo-EM structure [32] (see fig. 2e). On the other hand, there is no deformation when the hinge is bound (see fig. 2f). Very surprisingly, out of 299 analysed molecules, 149 were bound by the hinge and 150 bound by the heads; when both were bound, we considered the molecule both heads and hinge bound. This result demonstrates unexpectedly similar relative binding affinities to dsDNA of heads and hinge, broadly in line with the bulk EMSA and FP assays. Finally, we also observe both intra- and inter-molecular bridging, whereby two segments of DNA belonging to different molecules are simultaneously bound by the heads and hinge, in line with the transient cross-linking hypothesis proposed in the previous section (fig. 2h-i).

In summary, we discovered that yeast condensin can bind dsDNA through its hinge domain as well as through its heads, and that the two binding affinities are similar. Whilst the thermodynamics of binding may be similar, we argue that the kinetics of binding/unbinding may be very different for the two domains. We hypothesise that the “safety-belt” anchoring mechanism at the YcgI-BrnI domain may be very stable [32] (small *k*_*off*_ and small *k*_*on*_), whilst the kinetics at the hinge may be faster (large *k*_*off*_ and large *k*_*on*_). This reasoning could explain both the strong structural evidence for the seat-belt anchoring [32] and elusive DNA-hinge interaction as well as the potential for forming transient inter-molecular bridges, loops and cross-links.

### Coarse grained MD simulations of “sticky” SMCs recapitulate observed rheology

Having observed that the hinge is proficient in binding dsDNA we now aim to understand its consequences within the context of physiologically dense and entangled solutions of DNA. To this end we performed Molecular Dynamics (MD) simulations of entangled linear DNA under the action of SMCs displaying “sticky” patches that can bind to DNA polymer beads (see fig. 3a-b). Briefly, we modelled DNA molecules as Kremer-Grest bead-spring polymers [34] at fixed density, corresponding to 5% volume fraction. After equilibrating the system, we randomly loaded SMCs onto the polymers and allowed them to form both DNA loops *in cis* and inter-molecular bridges (Fig. 3a-c). Our SMC model is different from most paradigms in the literature [2, 24, 27, 38, 43–45] as we allow them to do both, form intra/inter molecular bridges through their patches and also form loops through extrusion whilst preserving the polymer topology (see SI for full details).

**FIG. 3.**
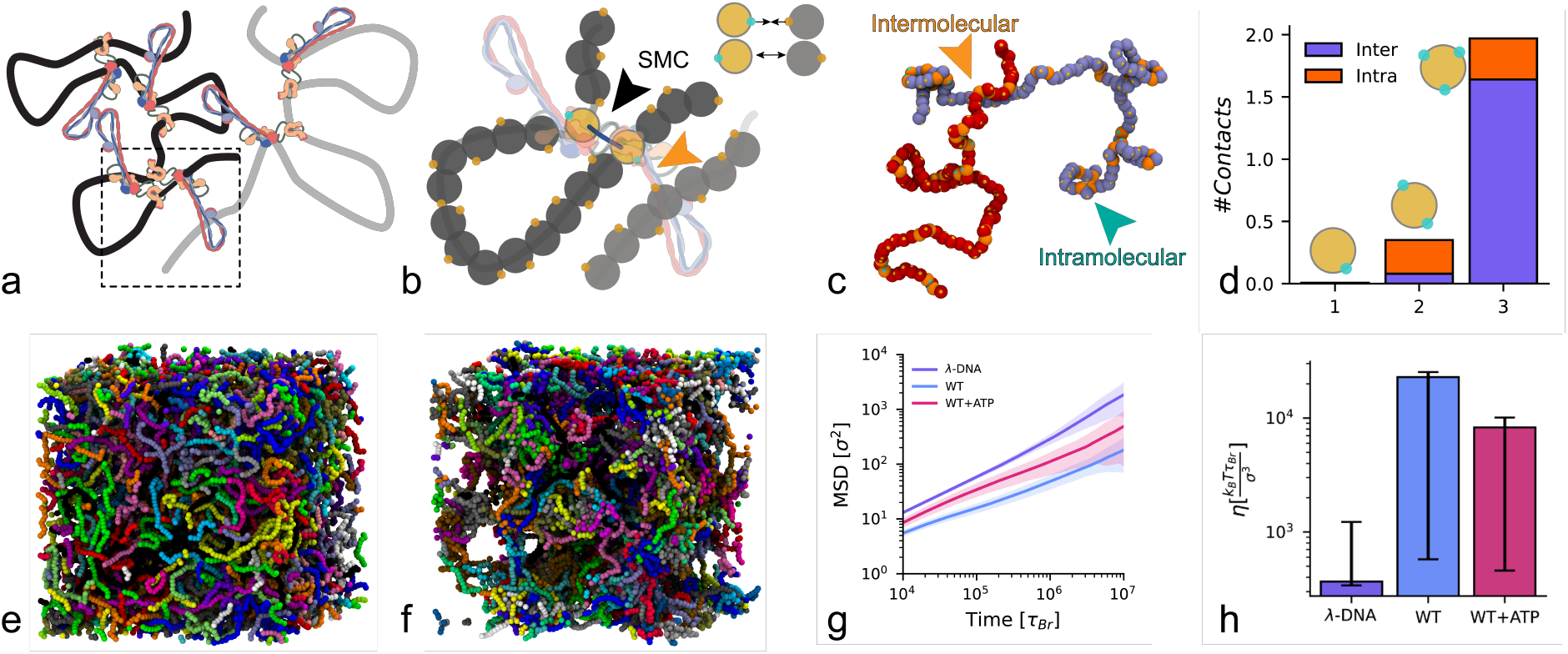
MD simulations of sticky SMCs recapitulate the thickening. **a**. Sketch of DNA with loops formed by SMCs. **b**. Bead-spring polymer modelling of the region in the dashed box showing the correspondence between patches (cyan) and hinge domain. **c**. Snapshot from simulations, highlighting intramolecular and intermolecular interactions stabilised by the patches. **d** Average number of contacts as a function of the number of patches on the beads. **e-f**. Snapshots of the simulation box in two cases: (**e**) in equilibrium with no SMC and (**f**) after loading 50 stikcy SMCs per polymer and allowing them to extrude loops. **g**. Average Mean Squared Displacement (MSD) of the polymers’ center of mass (standard deviation shaded) for the control case (*λ*-DNA) compared with the cases with SMC but no extrusion (WT) and the case with SMC allowed to extrude loops (WT+ATP). **h**. Viscosity computed from the stress-relaxation function (see SI) for the three cases in **g**. Notice that with ATP, the system is more fluid, in line with experiments.

To account for the formation of SMC clusters [19] we also explored the effect of having multiple DNA binding sites on each SMC bead: on average, less than one contact per SMC complex was seen for patches per SMC bead, while 2 contacts (mostly inter-molecular) were seen for *N*_*p*_ = 3 patches per SMC bead (fig. 3d). This implies that only one third of all SMC patches were bound to DNA at any one time and corresponds to the case in which there are two clustered, or stacked, SMCs per loop. In this scenario, the two SMCs are bound to the same DNA by their heads, and have two “free” hinges that can make inter or intra molecular contacts with other DNA segments (see fig. 3a-b).

We observed that once the SMCs are loaded, the system qualitatively displayed remarkable clustering (see snapshots fig. 3e-f). By computing the mean squared displacement of the centre of mass of the polymers, i.e. *δ*^2^*r*(*t*) = ⟨ [***r***_*CM*_ (*t* + *t*_0_) − ***r***_*CM*_ (*t*)]^2^⟩, we discovered that they also displayed a significantly slower dynamics, and large effective viscosity (fig. 3g-h). Interestingly, the systems with fewer patches (*N*_*p*_ = 0 or 1) displayed significant faster dynamics than the control (no SMC), due to the bottle-brush-like conformations assumed by the polymers [24, 27] (see SI).

To rationalise the experiments we thus placed on average 5 SMCs on each polymer (same stoichiometry as in the experiments) and compared simulations where SMCs were only bound to DNA (same as no ATP in experiments), and the case where SMCs were allowed to loop extrude (with ATP in epxeriments). The MSDs diplayed in Fig. 3g show that the dynamics of the polymers in presence of extrusion is faster than the no-extrusion case, as seen in experiments. Additionally, we computed the stress relaxation function *G*(*t*) through the autocorrelation of the out-of-diagonal components of the stress tensor [37, 46] and computed the zero-shear viscosity. In line with the results so far, the viscosity of the system appears larger for the case in which the SMCs are allowed to form loops before the dynamics is evaluated.

Overall, our simulations reveal that static SMCs can stabilise dynamic cross-links which render dense solutions of DNA more viscous. At the same time, allowing the SMCs to form loops lead to bottle-brush-like polymers [2, 38] which lowers the effective viscosity of the system by increasing the entropic cost of stabilising intermolecular interactions [24, 27]. Despite of this additional entropic repulsion, even in the presence of ATP we observe that SMCs have the net effect of increasing the viscoelasticity of the solution.

## DISCUSSION AND CONCLUSIONS

In summary, in this paper we have provided experimental and computational evidence that SMCs, and specifically yeast condensin, can stabilise inter-molecular interactions in solutions of dense DNA. To the best of our knowledge, this is the first time such evidence is presented. Most of the current models involving SMCs posit that they perform intra-chain loop extrusion. We reasoned that if this were the case, we would observe a speed up of the dynamics (or thinning of the rheology) of entangled DNA solutions (fig. 1a). On the contrary, we consistently observed that adding yeast condensin to a solution of entangled DNA drives a slowing down and thickening of the solution viscoelasticty (fig. 1e-g). This effect cannot be attributed to the mere presence of additional protein in the solution because (i) we observe increase in elasticity, implying the formation of DNA crosslinks and (ii) we observed the opposite behaviour with different proteins [40].

To understand the mechanisms underlying this change in rheology, we used biochemical assays and AFM to uncover that the hinge domain of yeast condensin is a proficient dsDNA binding sites (fig. 2c-d). Strikingly, we observed that it binds as strongly as its heads, which are very well-known DNA binding sites from structural studies [32] (fig. 2e-g). We further showed that SMCs can simultaneously bind dsDNA through their heads and hinge domains both intra- and inter-molecularly (fig. 2h-i).

To connect the rheology and single-molecule observations, we concluded this work by performing MD simulations and showing that modelling SMCs as “sticky” proteins that can both, stabilise dynamic intermolecular cross-links and form loops, we can fully recapitulate the rheology measurements (fig/ 3a-b,g-h). We therefore argue that in our *in vitro* experiments, SMCs do not exclusively form intrachain loops, but also stabilise intermolecular dynamic crosslinks therefore affecting the system’s rheology (fig. 4).

**FIG. 4.**
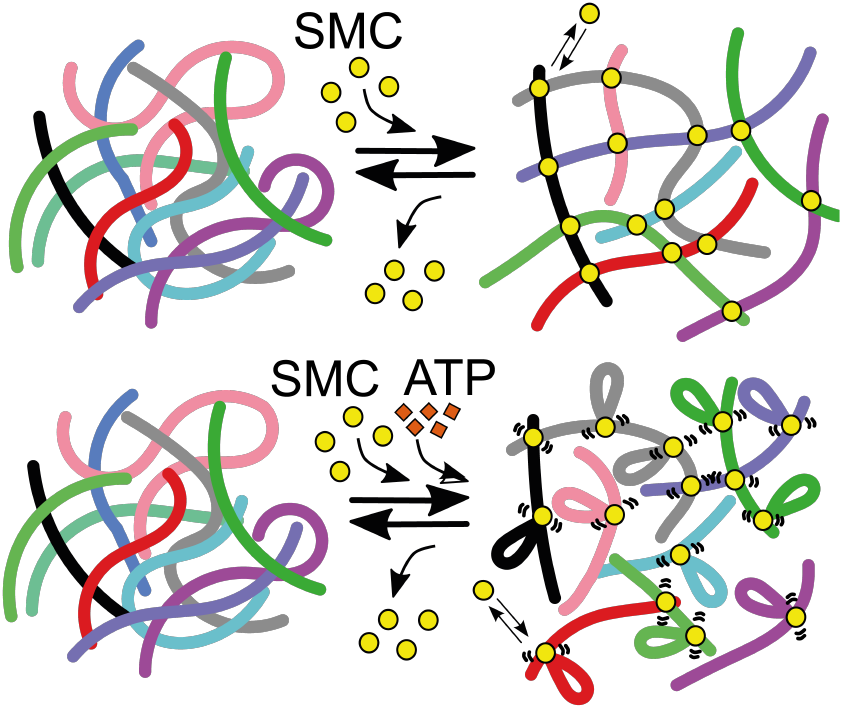
Modelling SMC’s action on entangled DNA. (top) SMCs loaded on DNA can form dynamic cross-links by simultaneously bridging dsDNA molecules with their heads and hinge. (bottom) The presence of ATP allows loop formation that competes with intermolecular bridging and lowers the viscosity of the system.

To our knowledge, this is the first time that the rheological impact of SMCs could be precisely quantified on a physiologically dense solution of DNA. Most of the current *in vitro* assays for loop extrusion [14] and loop capture [47] are performed on either single tethered DNA molecules or in dilute conditions. Both are rather distant from the conditions of density and entanglements experienced by DNA and chromatin *in vivo*. We thus argue that the effects uncovered in this work may be physiologically relevant and could in fact explain the puzzling observations from single molecule tracking *in vivo*, whereby deletion of cohesin typically induces a speed up of chromatin dynamics [23, 48]. To conclude, we argue that SMCs’s role in regulating chromatin dynamics may be more multifaceted and complex than previously thought. Their ability to form intermolecular DNA-DNA interactions should be considered in future models attempting to rationalise both chromosome conformation and dynamics. Even more important may be the interaction of SMCs with other proteins such as topoisomerase [36, 49]. Ultimately, measuring chromatin viscoelasticity in situ will reveal the impact of SMCs on genome viscoelasticity.

## ACKNOWLEDGEMENTS

DM acknowledges the Royal Society and the European Research Council (grant agreement No 947918, TAP) for funding. The authors also acknowledge the contribution of the COST Action Eutopia, CA17139. For the purpose of open access, the author has applied a Creative Commons Attribution (CC BY) licence to any Author Accepted Manuscript version arising from this submission. The authors thank Markus Hassler and Christian Häring for comments and feedback on the manuscript.

